# Hand-selective visual regions represent how to grasp 3D tools: brain decoding during real actions

**DOI:** 10.1101/2020.10.14.339606

**Authors:** Ethan Knights, Courtney Mansfield, Diana Tonin, Janak Saada, Fraser W. Smith, Stéphanie Rossit

## Abstract

Most neuroimaging experiments that investigate how tools and their actions are represented in the brain use visual paradigms where tools or hands are displayed as 2D images and no real movements are performed. These studies discovered selective visual responses in occipito-temporal and parietal cortices for viewing pictures of hands or tools, which are assumed to reflect action processing, but this has rarely been directly investigated. Here, we examined the responses of independently visually defined category-selective brain areas when participants grasped 3D tools. Using real action fMRI and multi-voxel pattern analysis, we found that grasp typicality representations (i.e., whether a tool is being grasped appropriately for use) were decodable from hand-selective areas in occipito-temporal and parietal cortices, but not from tool-, object-, or body-selective areas, even if partially overlapping. Importantly, these effects were exclusive for actions with tools, but not for biomechanically matched actions with control non-tools. In addition, decoding of grasp typicality was significantly higher in hand than tool-selective parietal regions. Notably, grasp typicality representations were automatically evoked even when there was no requirement for tool use and participants were naïve to object category (tool vs non-tools). Finding a specificity for typical tool grasping in hand-, rather than tool-, selective regions challenges the long-standing assumption that brain activation for viewing tool images reflects sensorimotor processing linked to tool manipulation. Instead our results show that typicality representations for tool grasping are automatically evoked in visual regions specialised for representing the human hand, the brain’s primary *tool* for interacting with the world.

**Significance Statement:** The unique ability of humans to manufacture and use tools is unsurpassed across the animal kingdom, with tool use considered a defining feature of our species. Most neuroscientific studies that investigate the brain mechanisms that support tool use, record brain activity while people simply view images of tools or hands and not when people perform actual hand movements with tools. Here we show that specific areas of the human visual system that preferentially process hands automatically encode how to appropriately grasp 3D tools, even when no actual tool use is required. These findings suggest that visual areas optimized for processing hands represent fundamental aspects of tool grasping in humans, such as which side they should be grasped for correct manipulation.

## INTRODUCTION

The emergence of handheld tools (e.g., a spoon) marks the beginning of a major discontinuity between humans and our closest primate relatives (Ambrose, 2001). Unlike other manipulable objects (e.g., books), tools are tightly associated with predictable motor routines (Johnson-Frey, 2004). A highly replicable functional imaging finding is that simply viewing tool pictures activates sensorimotor brain areas (Lewis, 2006), but what drives this functional selectivity? One popular idea is that this visually-evoked activation reflects the automatic extraction of information about the actions tools afford, like the hand movements required for their use (e.g., Martin et al., 1996; Fang & He, 2005). Similarly, tool-selective visual responses in Supramarginal (SMG) or posterior Middle Temporal Gyri (pMTG) are often interpreted as indirect evidence that these regions are involved in real tool manipulation (e.g., Buxbaum et al., 2006; Bach et al., 2010). Nevertheless, we would never grasp a picture of a tool and, more importantly, finding spatially overlapping activation between two tasks does not directly imply that the same neural representations are being triggered (Dinstein et al., 2008; Martin, 2016). In fact, intraparietal activation for viewing tool pictures vs grasping shows poor correspondence (Valyear et al., 2007; Gallivan et al., 2013), questioning the long-standing assumption that visual tool-selectivity represents sensorimotor aspects of manipulation.

Curiously, the visual regions activated by viewing pictures of hands in the left IPS (IPS-Hand) and Lateral Occipital Temporal Cortex (LOTC-Hand) overlap with their respective tool-selective areas (IPS-Tool; LOTC-Tool; Bracci et al., 2012; 2013; 2016). Stimulus features often described to drive the organisation of category-selective areas, like form (Coggan et al., 2016), animacy (Konkle & Caramazza, 2013) or manipulability (Mahon et al., 2007) poorly explain this shared topography because hands and tools differ on these dimensions. Instead, their overlap is suggested to result from a joint representation of high-level action information related to skilful object manipulation (Bracci et al., 2012; 2016; Striem-Amit et al., 2017), perhaps coding the function of hand configurations (Perini et al., 2014; Bracci et al., 2018). Arguably, the only way to directly test whether tool- or hand-selective visual areas carry information about tool actions is to examine their responses during real 3D tool manipulation. Yet, very few fMRI studies involve real tool manipulation (e.g., Gallivan, et al., 2009; Valyear et al., 2012; Brandi et al., 2014; Styrkowiec et al., 2019). To date, only Gallivan et al. (2013) investigated real tool manipulation in visually defined tool-selective regions and showed that IPS-/LOTC-Tool are indeed sensitive to coarsely different biomechanical actions (reaching vs grasping) with a pair of tongs. However, it remains unknown whether hand-selective visual areas represent properties of real hand movements with 3D tools, like the way they are typically grasped for subsequent use.

Here, an fMRI experiment involving real hand actions (Fig. 1) tested if visually defined hand- and tool-selective areas represented how to typically grasp 3D tools. Specifically, participants grasped 3D-printed tools in ways either consistent with their use (typical: by their handle) or not (atypical: by their functional-end; e.g., knife blade). As a control, non-tool bars (matched with the tools for elongation, width and depth; adapted from Brandi et al., 2014) were also grasped on their right or left sides to match as much as possible any biomechanical differences between typical and atypical actions. Multivoxel Pattern Analysis (MVPA) was used to assess whether different tool grasps (typical vs atypical) and non-tool grasps (right vs left), could be decoded from fMRI activity patterns within independent visually defined Regions of Interest (ROIs). Greater-than-chance decoding accuracy of typical vs atypical actions for tools, but not control non-tools, was interpreted as evidence that an area contains high-level typicality representations about how a tool should be grasped correctly for use (i.e., by its handle). This pattern of findings was expected only for the tool- and hand-selective areas since these are thought to support tool manipulation (e.g., Mahon & Caramazza, 2009; Striem-Amit et al., 2017).

**Figure 1.**
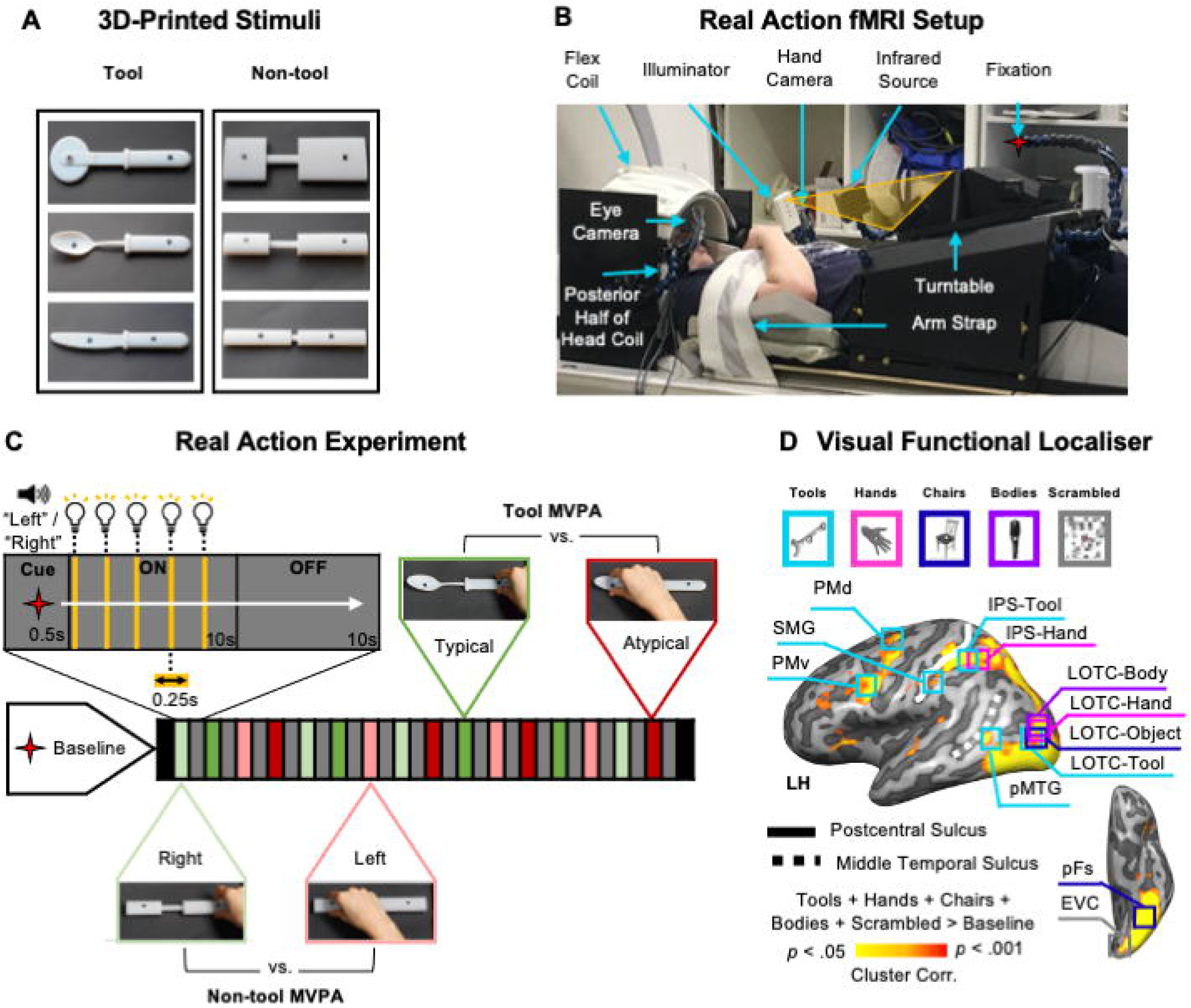
Experimental set-up and design. (A) 3D-printed tool and non-tool control object pairs (black markers on objects indicate grasp points) which were matched for elongation, width and depth such that tool and non-tool actions were biomechanically similar. (B) Side view of real action participant set-up used to present 3D objects at grasping distance (without the use of mirrors). Red star indicates fixation LED. The hand is shown at its starting position. (C) Timing and grasping tasks from subject’s point of view. During the 10s ON-block the object was illuminated 5 times cueing the participant to grasp the object each time by its left or right side (as per preceding auditory cue) with the right hand. Exemplar videos of trial types can be assessed here: https://osf.io/gsmyw/. This was followed by a 10s OFF-block involving no stimulation where the workspace remained dark. For MVPA, we treated tool and non-tool trials independently, where for the tools only, right- and left-sided grasps were typical and atypical grasps respectively (based on handle orientation). (D) Visual functional localizer experiment tools, hands, chairs, bodies and scrambled exemplar stimuli. For each individual participant independent ROIs were defined for MVPA (Table 1). The representative ROI locations are displayed on a group activation contrast map from the visual localizer (All conditions > (Baseline*5)) projected onto a left hemisphere cortical surface reconstruction of a reference brain (COLIN27 Talairach) available from the neuroElf package (http://neuroelf.net).

## Materials and Methods

### Participants

Twenty healthy participants (11 males) completed the real action fMRI experiment followed by a visual localizer experiment on a separate day. Data from one participant (male) was excluded from statistical analysis due to excessive head movements during the real action experiment (i.e., translation and rotation exceeded 1.5mm and 1.5° rotation) leaving a total sample of 19 participants (mean age = 23 years ± 4.2 years; age range = 18 - 34). All participants had normal or corrected-to-normal vision, no history of neurological or psychiatric disorders, were right-handed (Oldfield, 1971) and gave written consent in line with procedures approved by the School of Psychology ethics committee at the University of East Anglia.

### Real action 3D stimuli

Tool and non-tool object categories were designed (Autodesk Inc.) and 3D-printed (Objet30 Desktop) in VeroWhite material (Statasys): three common kitchen tools (knife, spoon and pizzacutter) and three non-tool control bars (see Fig.1A). Objects were secured to slots placed onto black pedestals used for stimulus presentation. Tools had identical handles (length x width x depth dimensions of 11.6cm x 1.9cm x 1.1cm) with different functional-ends attached (knife = 10.1cm x 1.9cm x 0.2cm; spoon = 10.1cm x 4.1cm x 0.7cm; pizzacutter = 10.1cm x 7.5cm x 0.2cm). To avoid motor or visual confounds, tools and non-tool pairs were carefully matched in terms of visual properties and kinematic requirements as much as possible. Specifically, non-tools were comprised of three cylindrical shapes (adapted from Brandi et al., 2014) with handle, neck and functional-end dimensions matched to each tool they controlled for, ensuring that grip size was matched between tool and non-tool pairs. In addition, all objects had small black stickers placed at pre-specified locations to indicate grasp points, ensuring that grasp position/reach-distance were identical between tool and non-tool pairs regardless of side to be grasped. To avoid familiarity confounds between tools and non-tool control stimuli we chose to use bars instead of scrambled tools and thus, our control non-tools were familiar, but had no specific associated function. Furthermore, each tool and non-tool pair were carefully matched for elongation so that any differences between conditions could not be explain by low-level shape preferences (e.g., Sakuraba et al., 2012; Brandi et al., 2014).

### Real action setup and apparatus

Participants were scanned in complete darkness using a head-tilted configuration that allowed direct viewing of the workspace and 3D stimuli without the use of mirrors (Fig. 1B) by tilting the head coil ~20° and padding the underside of each participants heads with foam cushions (NoMoCo Pillow, La Jolla, CA, USA). Objects were placed by an experimenter on a turntable above the participant’s pelvis and were only visible when illuminated (e.g., Fernández-Espejo et al., 2015; Fig. 1B). All stimuli were mounted such that they were aligned with participants’ midlines, never changed position while visible and were tilted away from the horizontal at an angle (~15°) to maximize visibility and grasp comfort. For stimulus presentation, the workspace and object were illuminated from the front using a bright white Light Emitting Diode (LED) attached to a flexible plastic stalk (Loc-line, Lockwood Products; Fig. 1B). To control for eye movements, participants were instructed to fixate a small red LED positioned above and behind objects such that they appeared in the lower visual field (Rossit et al., 2013). Throughout the experiment, participants’ right eye and arm movements were monitored online and recorded using two MR-compatible infrared-sensitive cameras (MRC Systems GmbH) to verify that participants performed the correct grasping movement (hand camera positioned over the left shoulder; Fig. 1B) and maintained fixation (eye camera beside the right eye; Fig. 1B). The likelihood of motion artefacts related to grasping was reduced by restraining the upper-right arm and providing support with additional cushions so that movements were performed by flexion around the elbow only (Culham, 2006). Auditory instructions were delivered to the participants through Earphones (Sensimetrics MRI-Compatible Insert Earphones Model S14, USA). At the beginning of the real action session, participant setup involved adjusting the exact position of: 1) stimuli and the hand to ensure reachability (average grasping distance between the “home” position and object = 43cm), 2) the illuminator to equally light all objects, 3) the fixation LED to meet the natural line of gaze (average distance from fixation to bridge nose = 91cm; visual angle = ~20°) and 4) the infrared-sensitive eye and hand cameras to monitor eye and hand movement errors. The experiment was controlled by a Matlab script (The MathWorks, USA R2010a) using the Psychophysics Toolbox (Brainard, 1997).

### Real action experimental paradigm

We used a powerful block-design fMRI paradigm, that maximised the contrast-to-noise ratio to generate a reliable estimate of the average response pattern (Mur et al., 2009) and improved detection of blood oxygenation level-dependent (BOLD) signal changes without significant interference from artefacts during overt movement (Birn et al., 2004). A block began with an auditory instruction (‘Left’ or ‘Right’; 0.5s) specifying which side of the upcoming object to grasp (Fig. 1C). During the ON-block (10s), the object was briefly illuminated for 0.25s five consecutive times (within 2s intervals) cueing the participant to grasp with a right-handed precision grip (i.e., index finger and thumb) along the vertical axis. Between actions, participants returned their hand to a “home” position with their right hand closed in a fist on their chest (see Fig. 1B). This brief object flashing presentation cycle during ON-blocks has been shown to maximise the signal-to-noise ratio in previous perceptual decoding experiments (Kay et al., 2008; Smith & Muckli, 2010) and eliminates the sensory confound from viewing hand movements (Rossit et al., 2013; Monaco et al., 2015). An OFF-block (10s) followed the stimulation block where the workspace remained dark and the experimenter placed the next stimulus. A single fMRI run included 16 blocks involving the four grasping conditions (i.e., typical tool, atypical tool, right non-tool and left non-tool) each with three repetitions (one per exemplar; every object was presented twice and grasped on each side once). An additional tool (whisk) and a non-tool object were presented on the remaining four blocks per run, but not analysed as they were not matched in dimensions due to a technical problem (the original control non-tool for the whisk was too large to allow rotation of the turntable within the scanner bore). On average participants completed six runs (minimum five, maximum seven) for a total of 18 repetitions per grasping condition. Block orders were pseudorandomised such that conditions were never repeated (two-back) and were preceded an equal amount of times by other conditions. Each functional scan lasted 356s, inclusive of start / end baseline fixation periods (14s). Each experimental session lasted ~2.25 hours (including setup, task practice and anatomical scan). Prior to the fMRI experiment, participants were familiarised with the setup and practiced the grasping tasks in a separate lab session (30 minutes) outside of the scanner. The hand and eye movement videos were monitored online and offline to exclude error trials from fMRI analysis. Two runs (of two separate participants) from the entire experiment were excluded from further analysis. In one of these blocks the participant failed to follow the grasping task instructions correctly (i.e., performing alternated left and right grasps) and for the remaining block another participant did not maintain fixation (i.e., made downward saccades toward objects). In the remaining runs that were analysed, participants made performance errors in <1% of experimental trials. The types of errors included: not reaching (3 trials, 2 participants), reaching in the wrong direction (1 trial, 1 participant) and downward eye saccades (5 trials, 3 participants). A one-way repeated measures ANOVA with 12 levels (i.e., the six exemplars across both left vs right grasping conditions) showed that the percentage of errors were equally distributed amongst trial types regardless of whether the percentage of hand and eye errors were combined or treated separately (all p’s > 0.28).

Crucially, since the tools’ handles were always oriented rightward, the right and left tool trials involved grasping tools either by their handle (typical) or functional-end (atypical), respectively. On the other hand, grasping non-tools did not involve a typical manipulation but only differed in grasp direction with right vs left grasps (Fig. 1C). We chose to present rightward oriented tool handles only, rather than alternate object orientation randomly between trials, to reduce total trial numbers (scanning times was already quite extensive with set-up) and due to technical limitations (i.e., the turntable’s rotation direction was fixed and it was difficult for the experimenter to manipulate tool orientation in the dark). Nevertheless, by comparing the decoding accuracies for each region between tool and non-tool grasps (which were matched for biomechanics) we ruled out the possibility that our typically manipulation simply reflected grasp direction. Specifically, we took the conservative approach that for an area to be sensitive to tool grasping typicality, it should not only show greater-than-chance decoding for typical vs atypical actions with tools (i.e. typicality), but also that the typicality decoding accuracy should be significantly greater than accuracy for biomechanically matched actions with our control non-tools (i.e. right vs left actions with non-tools).

### Visual Localizer

On a separate day from the real action experiment, participants completed a Bodies, Chairs, Tools and Hands (BOTH) visual localizer (adapted from Bracci et al., 2012; 2013; 2016) using a standard coil configuration (see MRI acquisition for details). Two sets of exemplar images were selected from previous stimuli databases (Bracci et al., 2012; 2013; 2016) that were chosen to match, as much as possible, the characteristics within the tool (i.e., identity & orientation), body (i.e., gender, body position & amount of skin shown), hand (i.e., position & orientation) and chair (i.e., materials, type & style) categories. Using a mirror attached to the head coil, participants viewed separate blocks (14s) of 14 different grayscale 2D pictures from a given category (400 x 400 pixels; 0.5s). Blank intervals separated individual stimuli (0.5s) and scrambled image blocks separated cycles of the four randomised category blocks. Throughout, participants fixated a superimposed bullseye on the centre of each image and, to encourage attention, performed a one-back repetition detection task where they made a right-handed button press whenever 2 successive photographs were identical. The 2D images stimuli were presented with an LCD projector (SilentVision SV-6011 LCD, Avotech Inc.). A single fMRI run included 24 category blocks (6 reps per condition) with blank fixation baseline periods (14s) at the beginning and the end of the experiment. Each localizer scan lasted 448s and, on average, participants completed 4 runs (minimum 3, maximum 4) for a total of 24 reps per condition. The entire localizer session lasted ~50 minutes after including the time taken to acquire a high-resolution anatomical scan and setup participants.

### MRI Acquisition

The BOLD fMRI measurements were acquired using a 3T wide bore GE-750 Discovery MR scanner at the Norfolk & Norwich University Hospital (Norwich, UK). To achieve a good signal to noise ratio during the real action fMRI experiment, the posterior half of a 21-channel receive-only coil was tilted and a 16-channel receive-only flex coil was suspended over the anterior-superior part of the skull (see Fig. 1B). A T2*-weighted single-shot gradient Echo-Planer Imaging (EPI) sequence was used throughout the real action experiment to acquire 178 functional MRI volumes (Time to Repetition (TR) = 2000ms; Voxel Resolution (VR) = 3.3 x 3.3 x 3.3mm; Time to Echo (TE) = 30ms; Flip Angle (FA) = 78°; Field of View (FOV) = 211x 211mm; Matrix Size (MS) = 64 x 64) that comprised 35 oblique slices (no gap) acquired at 30° with respect to AC-PC, to provide near whole brain coverage. A T1-weighted anatomical image with 196 slices was acquired at the beginning of the session using BRAVO sequences (TR = 2000ms; TE = 30ms; FOV = 230mm x 230mm x 230mm; FA = 9°; MS = 256 x 256; Voxel size = 0.9 x 0.9 x 0.9mm).

For visual localizer sessions, a full 21-channel head coil was used to obtain 224 functional MRI volumes (Time to Repetition (TR) = 2000ms; Voxel Resolution (VR) = 3.3 x 3.3 x 3.3mm; Time to Echo (TE) = 30ms; Flip Angle (FA) = 78°; Field of View (FOV) = 211x 211mm; Matrix Size (MS) = 64 x 64). A high resolution T1-weighted anatomical image with 196 slices was acquired before the localizer runs (TR = 2000ms; TE = 30ms; FOV = 230mm x 230mm x 230mm; FA = 9°; MS = 256 x 256; Voxel size = 0.9 x 0.9 x 0.9mm). Localizer datasets for two participants were retrieved from another study from our group (Rossit et al., 2018) where the identical paradigm was performed when acquiring volumes using a Siemens whole-body 3T MAGNETOM Prisma fit scanner with a 64-channel head coil and integrated parallel imaging techniques at the Scannexus imaging centre (Maastricht, The Netherlands) and comparable acquisition parameters (Functional scans: TR = 2000 ms; TE = 30 ms; FA = 77°; FOV = 216mm; matrix size = 72 x 72; Anatomical scan: T1-weighted anatomical image: TR = 2250ms; TE = 2.21ms; FA = 9°; FOV = 256 mm; matrix size = 256 x 256).

### Data Preprocessing

Preprocessing and ROI definitions were performed using BrainVoyager QX (version 2.8.2) (Brain Innovation, Maastricht, The Netherlands). BrainVoyager’s 3D motion correction (sinc interpolation) aligned each functional volume within a run to the functional volume acquired closest in time to the anatomical scan (e.g., Rossit et al., 2013). Slice scan time correction (ascending and interleaved) and high-pass temporal filtering (2 cycles/run) was also performed. Functional data were superimposed on to the anatomical brain images acquired during the localizer paradigm that were previously aligned to the plane of the anterior-posterior commissure and transformed into standard stereotaxic space (Talairach, & Tournoux, 1988). Excessive motion was screened by examining the time-course movies and motion plots created with the motion-correction algorithms for each run. No spatial smoothing was applied.

To estimate activity in the localizer experiment, a predictor was used per image condition (i.e., Bodies, Objects, Tools, Hands and Scrambled) in a single-subject general linear model (GLM). Predictors were created from boxcar functions that were convolved with a standard *2y* model of the hemodynamic response function (Boynton et al., 1996) and aligned to the onset of the stimulus with durations matching block length. The baseline epochs were excluded from the model, and therefore, all regression coefficients were defined relative to this baseline activity. This process was repeated for the real action experiment, using 16 separate predictors for each block of stimulation independently per run (12 exemplars - knife typical, knife atypical, spoon typical etc. plus 4 foil trials) and 6 motion regressors (confound predictors). These estimates (beta weights) from the real action experiment were used as the input to the pattern classifier.

### Visual Localizer Regions of interest (ROIs)

Twelve visual ROIs were defined at the individual participant level from the independent BOTH localizer data by drawing a cube (15 voxels^3^) around the peak of activity from previously reported volumetric contrasts (see list below; Fig. 1D; Table 1) set at a threshold of *p* < .005 (Gallivan et al., 2013) or, if no activity was identified, of *p* < .01 (Bracci et al., 2016). In cases where no activity was observed, the ROI was omitted for that participant (see Table 1). Given the predominantly left lateralised nature of tool-processing (Lewis, 2006), all individual participant ROIs were defined in the left hemisphere (Bracci et al., 2012; 2013; 2016; Peelen et al., 2013). Six tool-selective ROIs commonly described in left frontoparietal and occipitotemporal cortices were identified by contrasting activation for tool pictures vs other object pictures (IPS-Tool; SMG; dorsal and ventral Premotor Cortex (PMd; PMv), LOTC-Tool; pMTG; Martin et al., 1996; Grafton et al., 1997). Moreover, two hand-selective ROIs were identified in LOTC (LOTC-Hand) and IPS (IPS-Hand) by contrasting activation for hand pictures vs pictures of other body parts (Bracci et al., 2012; 2016; 2018; Peelen et al., 2013; Palser & Cavina-Pratesi, 2018). Additionally, we defined a body-selective (LOTC-Body; Bodies > Chairs; Bracci & de Beeck, 2016), two object-selective ROIs (LOTC-Object; posterior Fusiform, pFs; Chairs > Scrambled; Bracci & de Beeck, 2016; Hutchison et al., 2014) and an Early Visual Cortex ROI (EVC; All Categories > Baseline; Bracci & de Beeck, 2016). The ROI locations were verified by a senior author (S.R.) with respect to the following anatomical guidelines and contrasts:

- Lateral Occipitotemporal Cortex-Object selective (LOTC-Object) - (Chairs > Scrambled) (Hutchison et al., 2014; Bracci & de Beeck, 2016) – defined by selecting the peak of activation near the Lateral Occipital Sulcus (LOS; Hutchison et al., 2014; Bracci & de Beeck, 2016; Malach et al., 1995; Grill-Spector et al., 1999; 2001).
- Lateral Occipitotemporal Cortex-Body selective (LOTC-Body) - (Bodies > Chairs) (Bracci & de Beeck, 2016) – defined by selecting the peak of activation near the LOS and inferior to the left Extrastriate Body Area (EBA; Valyear & Culham, 2010) which was identified by the contrast ((Bodies + Hands) > Chairs) (adapted from Bracci, et al., 2010; ((Whole Bodies + Body Parts) > (Hands + Chairs))). EBA was not included in the analysis.
- Lateral Occipitotemporal Cortex-Hand selective (LOTC-Hand) - ((Hands > Chairs) AND (Hands > Bodies)) (Bracci & de Beeck, 2016) - defined by selecting the peak of activation near the LOS. These were often anterior to LOTC-Body (Bracci et al., 2010; 2016).
- Lateral Occipitotemporal Cortex-Tool selective (LOTC-Tool) - (Tools > Chairs) (Bracci, et al., 2012; Hutchison et al., 2014) - defined by selecting the peak of activation near the LOS. These often closely overlapped LOTC-Hand (Bracci, et al., 2012).
- Posterior Middle Temporal Gyrus (pMTG) - (Tools > Chairs) (Hutchison, et al., 2014; Valyear & Culham, 2010) - defined by selecting the peak of activation on the pMTG, more lateral, ventral and anterior to EBA (Hutchison et al., 2014). We selected the peak anterior to the Anterior Occipital Sulcus (AOS), as the MTG is in the temporal lobe and the AOS separates the temporal from the occipital (Damasio, 1995).
- Posterior Fusiform Sulcus (pFs) - (Chairs > Scrambled) (Hutchison, et al., 2014) - defined by selecting the peak of activation in the posterior aspect of the fusiform gyrus, extending into the occipitotemporal sulcus (Hutchinson, et al., 2014).
- Intraparietal Sulcus-Hand selective (IPS-Hand) - (Hands > Chairs) (Bracci, et al. 2016; Bracci & de Beeck, 2016) - defined by selecting the peak of activation on the IPS (Bracci & de Beeck, 2016).
- Intraparietal Sulcus-Tool selective (IPS-Tool) - (Tools > Scrambled) (Bracci, et al., 2016; Bracci et al., 2016) - defined by selecting the peak of activation on the IPS (Bracci & Op de Beeck, 2016).
- Supramarginal Gyrus (SMG) - (Tools > Scrambled) (Creem-Regehr, et al., 2007) - defined by selecting the peak of activation located most anterior along the SMG (Peeters, et al., 2013), lateral to the anterior segment of the IPS (Gallivan, et al., 2013), posterior to the Precentral Suclus (PreCS) and superior to the lateral sulcus (Ariani, et al., 2015).
- Dorsal Premotor Cortex (PMd) - (Tools > Scrambled) - defined by selecting the peak of activation at the junction of the PreCS and the superior frontal sulcus (Gallivan, 2013; Ariani, et al., 2015).
- Ventral Premotor Cortex (PMv) - (Tools > Scrambled) (Creem-Regehr, et al., 2007) - defined by selecting the voxels inferior and posterior to the junction between the inferior frontal sulcus and the PreCS (Gallivan et al., 2013).
- Early Visual Cortex (EVC) - (All Conditions > Baseline) (Bracci & de Beeck, 2016) - defined by selecting the voxels in the occipital cortex near the calcarine sulcus (Singhal, et al., 2013).

**Table 1.**
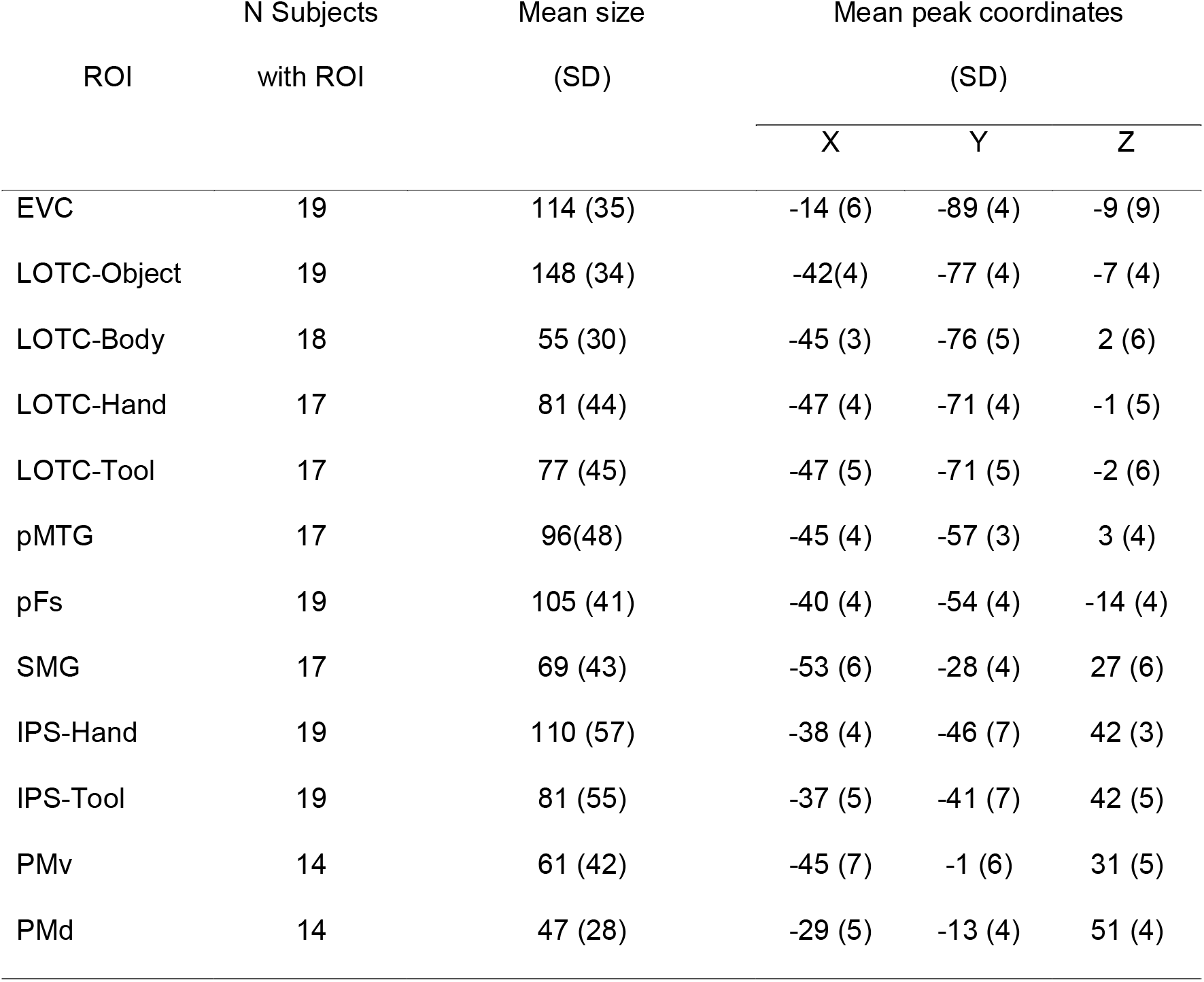
Visual Localizer ROI descriptives. ROI counts per subject with their mean sizes (voxels) and peak coordinates (Talairach).

### Pattern Classification

We performed MVPA independently for tool and non-tool trial types. Independent linear pattern classifiers (linear Support Vector Machine; SVM), were trained to learn the mapping between a set of brain-activity patterns (beta values computed from single blocks of activity) from the visual ROIs and the type of grasp being performed with the tools (typical vs atypical) or non-tools (right vs left). To test the performance of our classifiers, decoding accuracy was assessed using an n-fold leave on run out cross validation procedure; thus our models were built from n – 1 runs and were tested on the independent nth run (repeated for the n different possible partitions of runs in this scheme (Duda et al., 2001; Smith et al., 2010; 2015; Gallivan et al., 2016) before averaging across n iterations to produce a representative decoding accuracy measure per participant and per ROI. The activity of each ROI was normalised (separately for training and test data) within a range of −1 to +1 before input the SVM (Chang & Lin, 2011) and the linear SVM algorithm was implemented using the default parameters provided in the LibSVM toolbox (C = 1). Pattern classification was performed with a combination of in-house scripts (Smith et al., 2010; 2015) using Matlab with the Neuroelf toolbox (version 0.9c; http://neuroelf.net/) and a linear SVM classifier (libSVM 2.12 toolbox; https://csie.ntu.edu.tw/~cjlin/libsvm/).

### Statistical Analysis

One-tailed one-sample t-tests were used to test for above chance decoding for tool and non-tool action classifications in every ROI independently. If the pattern of results was consistent with our hypothesis (i.e., decoding accuracy was significantly above chance for tools, but not non-tools), we further ran a one-tailed pairwise t-tests to compare if decoding accuracy was significantly higher for tools than non-tools. Additionally, to test for differences in decoding accuracy between ROIs we used repeated measures 2 x 2 ANOVAs with ROI (tool vs hand selective) and object category (tool vs non-tool) as within-subject factors. Then, to test if univariate differences would differ between grasp types for the tools, but not non-tools we ran 2 x 2 ANOVAs with grasp type (typical/right vs atypical/left) and object category (tools vs non-tools) by entering mean beta weights for each ROI. Separately for each set of analyses we corrected for multiple comparisons with False Discovery Rate (FDR) correction of q ≤ 0.05 (Benjamini & Hochberg, 1995; Benjamini & Yekutieli, 2001) across the number of tests. Only significant results are reported (see Fig 2).

**Figure 2.**
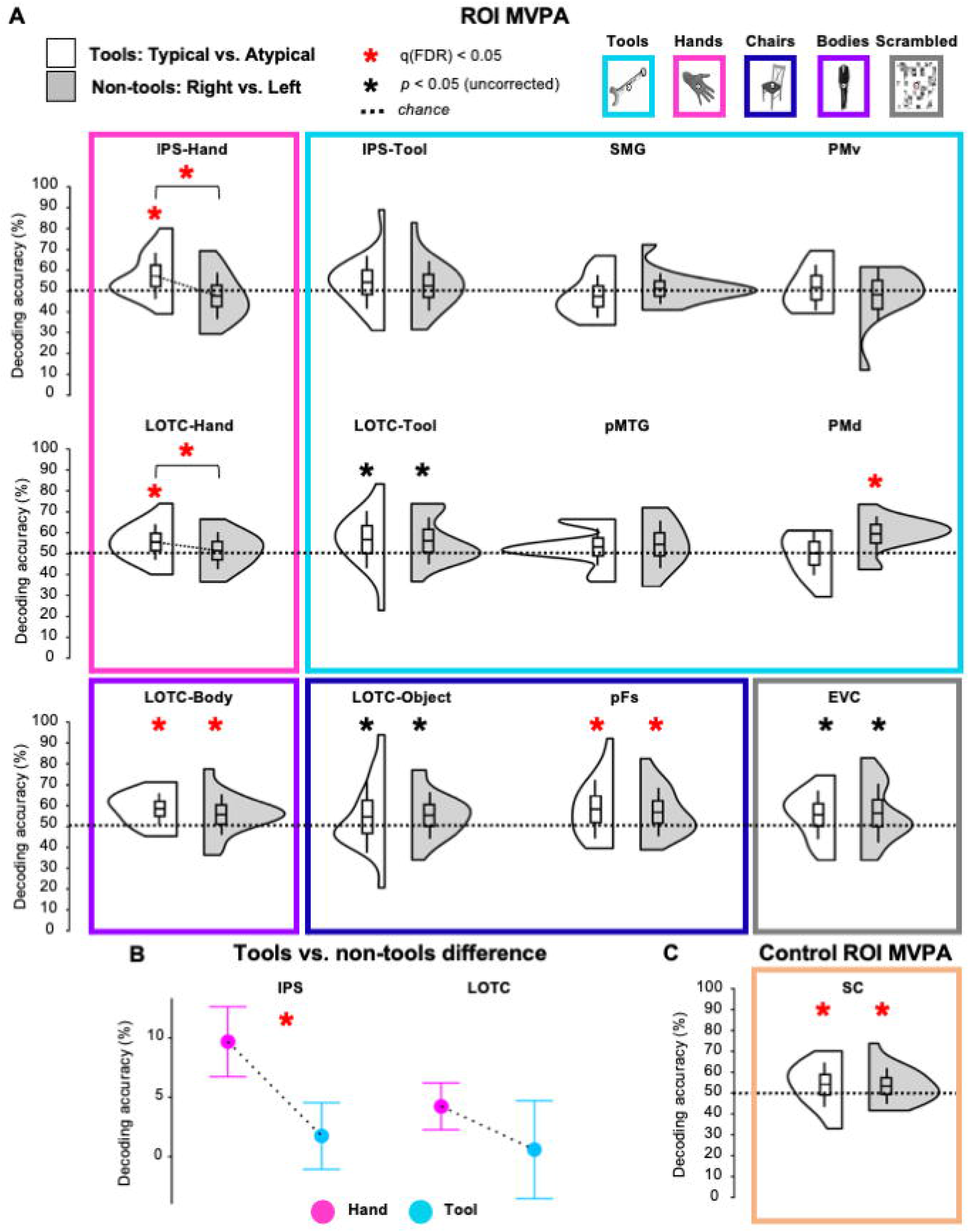
Grasp type decoding results in left hemisphere ROIs. (A) Violin plots of MVPA data from visual localizer ROIs for the typical vs atypical classification of grasping tools (white violins) and, non-tool control grasping (right vs left decoding; grey violins). Box plot centre lines are mean decoding accuracy while their edges and whiskers show ± 1 SD and ± 2 SEM, respectively. Decoding accuracies of typical vs atypical grasping in IPS and LOTC hand-selective cortex (pink) are significantly greater-than-chance for tools, but not non-tools. (B) ANOVA results comparing the difference of decoding accuracy between tools (typical vs atypical) and vs non-tools (right vs left) for the partially overlapping hand- and tool-selective ROIs within the IPS and LOTC. (C) Violin plot of MVPA data for control ROI in somatosensory cortex (SC) based on an independent contrast (all actions > baseline) from real action experiment showing significant decoding of grasp type for both tools and non-tools. Red asterisks show FDR-corrected results while black asterisks show uncorrected results.

To test for evidence for the null hypothesis over an alternative hypothesis, we supplemented null-hypothesis significance tests with Bayes factors (BF; Wagenmakers, 2007; Rouder et al., 2009). Bayes factors were estimated using the bayesFactor toolbox in Matlab (version 1.1; https://klabhub.github.io/bayesFactor/). The Jeffreys-Zellner-Siow default prior on effect sizes was used (Rouder, Morey, Speckman, & Province, 2012) and BF’s were interpreted according to criteria set out by Jeffreys (1961; cited from Jarosz & Wiley, 2014) where a BF_01_ between 1-3 and > 3 indicates ‘anecdotal’ and ‘substantial’ evidence in favour of the null, respectively.

### Data Availability

Stimuli, code for running experiment and for MVPA analyses and ROI data are accessible from Open Science Framework at: https://osf.io/zxnpv/. Full raw MRI dataset (real action and visual localizer) is accessible from OpenNEURO at: https://openneuro.org/datasets/ds003342/versions/1.0.0.

## Results

In line with our predictions, as can be seen in Fig. 2, a one-sample t-test against chance (50%) showed that SVM decoding accuracy (FDR-corrected) from hand-selective ROIs in LOTC and IPS were significantly greater-than-chance when discriminating typical vs atypical actions with tools (LOTC-Hand accuracy = 56% ± (SD) 0.9%, *t*(16) = 2.73, *p* = 0.007, *d* = 0.66; IPS-Hand accuracy = 57% ± 0.11%, *t*(18) = 2.72, *p* = 0.007, *d* = 0.62), but not biomechanically-matched actions with non-tools (right vs left; LOTC-Hand: *p* = 0.252, IPS-Hand: *p* = 0.844). In fact, there was substantial evidence in favour of null decoding of non-tool actions for the IPS ROI (LOTC-Hand: BF = 2.29; IPS-Hand = 8.4). Importantly, results from a stringent between-classification paired samples t-test also further supported this: typicality decoding accuracy from both LOTC-Hand and IPS-Hand was significantly higher for tools than for biomechanically-matched actions with non-tools (LOTC-Hand: *t*(16) = 2.11, *p* = 0.026, *d* = 0.51; IPS-Hand: *t*(18) = 3.26, *p* = 0.002, *d* = 0.75; Fig. 2A and Fig. 2B).

No other visual ROI, including tool-selective areas, displayed the same pattern of findings as hand-selective areas (Fig. 2A and Fig. 2B). For tool-selective ROIs, decoding accuracy was not significantly greater-than-chance for classifying actions with tools or non-tools (all p’s > 0.024), with the Bayesian approach demonstrating strong evidence in favour of the null for PMv (tool: BF_01_ = 3.23; non-tool: BF_01_ = 6.85) and SMG tool decoding (tool: BF_01_ = 8.85; other BF_01_‘s < 1.08). The exception to this was tool-selective PMd which was found to decode significantly above chance actions with non-tools (accuracy = 59% ± 0.08% *t*(13) = 4.11, *p* = 0.001, *d* = 1.1; Fig. 2A), but not tools (BF_01_ = 4.42). As for object- and body-selective areas, LOTC-Object decoding accuracy did not differ from chance for tools or non-tools (p > 0.026), though evidence in favour of the null was anecdotal (BF_01’s_ < 1.33), whereas pFs and LOTC-Body decoded actions above chance with both tools (pFs: accuracy = 58% ± 0.14% *t*(18) = 2.57, *p* = 0.01, *d* = 0.59; LOTC-Body: accuracy = 59% ± 0.08% (*t*(17) = 4.75, *p* < 0.001, *d* = 1.12) and non-tools (pFs: accuracy = 57% ± 0.12% *t*(18) = 2.59, *p* = 0.009, *d* = 0.59; LOTC-Body: accuracy = 56% ± 0.10% (*t*(17) = 2.46, *p* = 0.012, *d* = 0.58; Fig. 2A). Like many of the tool-selective ROIs, the control EVC ROI was not found to decode actions with either type of object (*p*’s < 0.026), albeit evidence in favour of the null was anedoctal (BF_01_’s > 0.37).

Since we obtained a different pattern of results for LOTC and IPS ROIs that were hand-vs tool-selective, we compared the decoding accuracies between these regions with a repeated measures ANOVA with ROI (hands vs tool-selective) and object category (tool vs non-tools) as within-factors. As shown in Fig. 2B, there was a significant interaction between ROI and object category in IPS (*F*(1,18) = 5.94, *p* = 0.025, η^2^ = 0.25). Post-hoc t-tests showed that for IPS-Hand, grasp type decoding was significantly higher for tools than non-tools (mean difference = 0.1%, SE = 0.03%; *p* = 0.004), but not for IPS-Tool (mean difference = 0.02%, SE = 0.03%). However, for LOTC this interaction was not significant (*p* = 0.379; Fig. 2B), nor were the remaining main effects (all *p*’s > 0.367).

Next, we examined whether significant decoding in hand-selective cortex could be accounted for by low-level sensory differences between the tools’ handles and functional-ends. First, to test the possibility that tool-specific decoding in hand-selective cortex could be driven by simple textural differences (e.g., a smooth handle vs a serrated knife blade), we repeated the analysis using a left somatosensory cortex ROI (SC; defined by selecting the peak voxel in the postcentral gyrus in the same subjects with an independent univariate contrast of All Grasps > Baseline; Fabbri et al., 2014, 2016). However, unlike the higher accuracies for grasping tools than non-tools in the hand-selective ROIs, grasp type decoding in SC was significantly greater-than-chance for both tool (accuracy = 57% ± 0.11%, *t*(18) = 3.04, *p* = 0.004, *d* = 0.7) and non-tools (accuracy = 57% ± 0.09% *t*(18) = 3.45, *p* = 0.001, *d* = 0.79; Fig. 1C). This indicates that tool-specific decoding in hand-selective cortex cannot be solely explained by somatosensory differences in the stimuli. Second, we tested if size differences between our objects, and thus grip size, could drive tool-specific decoding in hand-selective cortex (i.e., the functional-end of the tool being wider than its handle for the spoon and pizza cutter). However, decoding analysis of grip size (i.e., smaller vs larger objects) that collapsed across object category (i.e., tools and non-tools) was not significant for any visual ROI (all *p*’s ≥ 0.1; Fig. 3) and evidence in favour of the null was strong for most ROIs including IPS-Hand (BF_01_ = 8), EVC (BF_01_ = 3.22), LOTC-Object (BF01 = 4.93), pFs (BF01 = 5.97), SMG (BF01 = 3.33), PMv (BF01 = 3.91) and PMd (BF01 = 3.56; all other BF_01_’s > 1.84). Taken together, these findings suggest that hand-selective regions, particularly in the IPS, are sensitive to whether a tool is grasped correctly by its handle or not, and that these effects are not simply due to textural or size differences between the stimuli used or actions performed.

**Figure 3.**
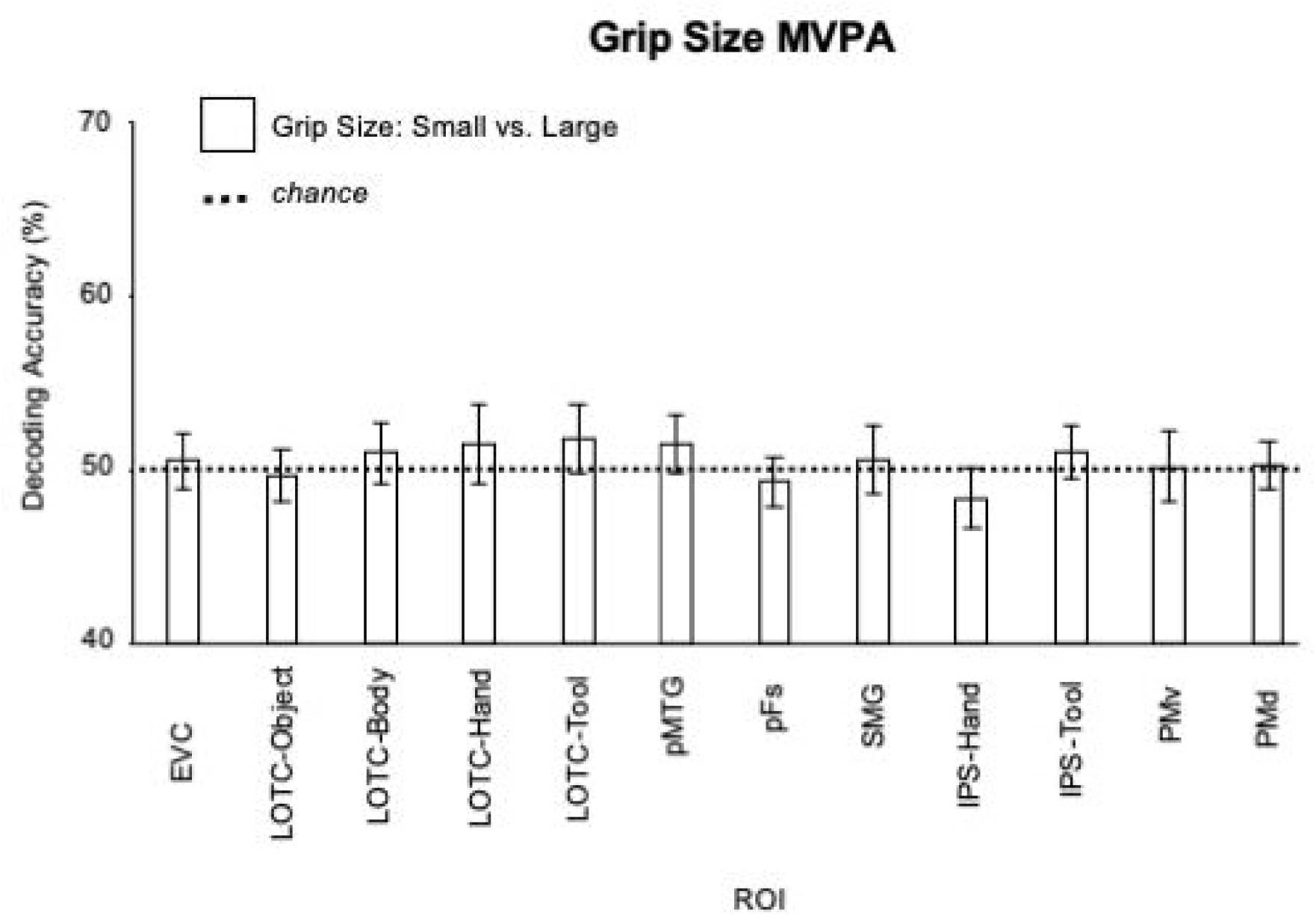
Mean activation (β) per ROI and condition used as input for pattern classification and univariate analyses. Error bars represent ± 1SEM.

In addition, we found that the significant decoding accuracies reported here do not simply reflect the overall response amplitudes within each ROI. When we analysed the mean beta weights in ANOVAs with grasp type and object category as within-subject factors for each ROI (i.e., as done in conventional univariate analysis; see Fig. 4), the only significant effect observed was a main effect of object category (unrelated to typicality), where greater activation was found for tools relative to non-tools in LOTC-Tool (F(1,16) = 9.25, p = 0.008, η^2^ = 0.37; mean difference = 0.1, SE = 0.03), pFs (F(1,18) = 8.68, p = 0.009, η^2^ = 0.33; mean difference = 0.07, SE = 0.02) and SMG (F(1,16) = 10.5, p = 0.005, η^2^ = 0.4; mean difference = 0.089, SE = 0.03).

**Figure 4.**
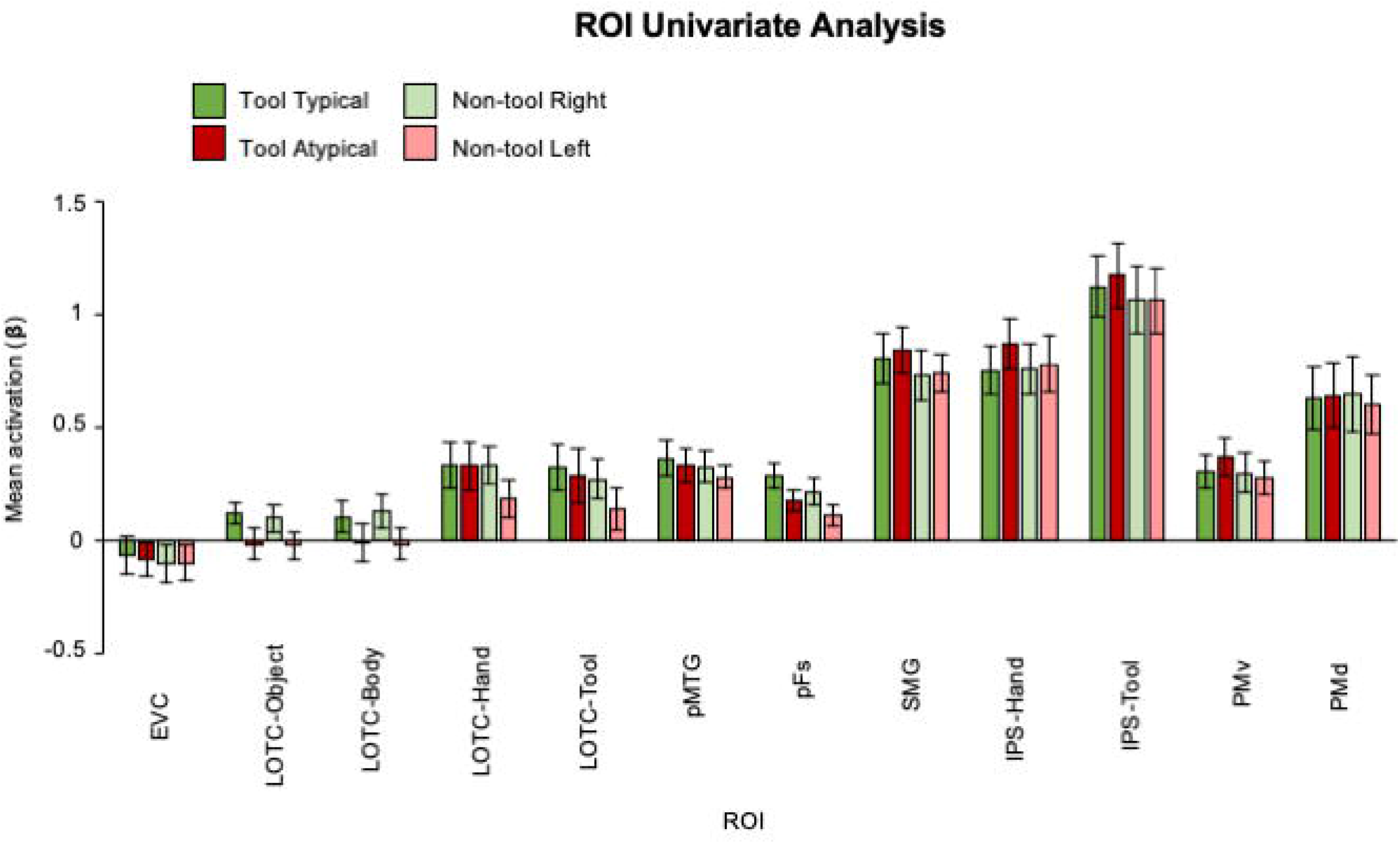
Grip size decoding results. Mean decoding accuracy in visual localizer ROIs and SC for the small versus large classification collapsed across object category. Error bars represent ±1 SEM.

## Discussion

Our understanding of how the human brain represents object properties (Kanwisher, 2010) and simple hand movements (Gallivan & Culham, 2015) has significantly advanced in last few decades, however far less is known about the neural representations that underpin real actions involving 3D tools (Valyear et al., 2017). Most neuroimaging experiments that investigate how tools and their associated actions are represented in the brain have used visual paradigms where objects and body-parts are displayed as 2D images (Ishibashi et al., 2016). These studies have discovered a tight anatomical and functional relationship between hand- and tool-selective areas in LOTC and IPS, thought to reflect action-related processing, however this was yet to be directly tested (Bracci et al., 2012; 2013; 2016; Peelen et al., 2013; Striem-Amit et al., 2017; Maimon Mor, 2020). Here we defined visually category-selective areas and investigated if they were sensitive to real action affordances involving 3D tools. We found the first evidence that hand-selective cortex (left IPS-Hand and LOTC-Hand) represents whether a 3D tool is being grasped appropriately by its handle. Remarkably, the same effects were not observed in tool-, object-, or body-selective areas, even when these areas overlapped with hand-selective voxels in both IPS and LOTC.

Our results indicate that visual hand-selective areas in parietal and occipital cortices process sensorimotor affordances of typicality for hand movements with 3D tools. Importantly, these action-related representations were detected exclusively for actions with tools, but not for biomechanically matched actions with non-tools. This tool-specificity was particularly evident in IPS-hand, where Bayesian evidence demonstrated that decoding of grasp type with non-tools was not possible. Our findings shed light into the features of sensorimotor processing in hand-selective areas. First, their representations are sensitive to concepts acquired through experience (i.e., knowing how to grasp tools appropriately is a learnt skill; Martin, 2007; Valyear et al., 2012), fitting with evidence showing that learning about how to manipulate tools (Weisberg et al., 2007) or even performing such actions (Valyear et al., 2012; Brandi et al., 2014; Styrkowiec et al., 2019) affects LOTC and IPS activity. Second, information processed by hand-selective cortex is represented in an abstract format beyond low level properties (e.g., basic kinematics), since Bayesian evidence strongly suggested that decoding grip size was not possible. This fits well with reports that hand-/tool-selective overlap exists in people born without vision (Peelen et al., 2013) or without hands (Striem-Amit et al., 2017) suggesting that their development is driven by similarities in how they process non-sensory tool information. In addition, our data also resonates with previous studies showing that tool-selective areas in pMTG/LOTC and IPS represent abstract action goals (reach vs grasp) regardless of biomechanics (Gallivan et al., 2013; Jacobs et al., 2010), albeit our findings were observed for hand-selective areas only. Third, our study shows that these high-level representations are automatically evoked (Valyear et al., 2007; Valyear et al., 2012; Grezes et al., 2003) as throughout the real-action fMRI task there was no explicit requirement to use the tools and participants were never told that we were investigating ‘tools’. Here we demonstrate that these principles, frequently described to support tool-use (Gibson, 1979; Imamizu et al., 2003; Maravita & Iriki 2004; Umilta et al., 2008; Lingnau & Downing, 2015), apply to brain areas specialised for representing the human hand, our primary *tool* for interacting with the world.

An intriguing aspect of our results is that typicality decoding was successful using activity patterns from hand-selective, but not overlapping parts of tool-selective cortex in the LOTC and IPS. Bayesian evidence only anecdotally supported the possibility that decoding was null from tool-selective areas, but significantly stronger typicality decoding was observed for IPS-Hand than IPS-Tool during tool, but not non-tool grasps. In contrast to previous picture viewing fMRI studies showing that overlapping hand- and tool-selective regions exhibit similar responses (Bracci et al., 2012; Bracci & Peelen, 2013; Bracci et al., 2016), our findings uniquely support previous speculations that hand-selective regions could be functionally distinct from tool-selective regions despite their anatomical overlap (Bracci et al., 2012; Striem-Amit et al., 2017). This pattern of results is unlikely to be driven by differences in ROI radius (Etzel et al., 2013) since voxel size differences were negligible between hand- and tool-selective ROIs (mean difference: IPS: 29; LOTC: 4). In fact, if category-related results were merely caused by ROI size, then significant decoding should have also been observed in the much larger LOTC-Object ROI (see Table 1). Alternatively, successful higher decoding in hand than tool-selective areas might reflect that our task simply required grasping-to-touch the tools, rather than their utilisation. That is, coding in category-selective areas might operate in an effector-dependent manner, akin to how tool-selective pMTG/LOTC codes the type of action being performed when holding a pair of tongs, but not if being performed by the hand alone (Gallivan et al., 2013). In line with this interpretation, neural representations in LOTC-Hand of one-handed amputees are also known to become richer as prosthetic usage increases (Van den Heiligenberg 2018), which, again, indicates that the representations in hand-selective cortex depend on effector use. An alternative, but not mutually exclusive, possibility is that only tool-use actions elicit tool-selective representations (see Randerath et al., 2010 for distinct lesion sites associated with grasping vs using tools) because of the cognitively taxing demands these complex actions rely on, such as retrieving knowledge about manipulation hierarchies (Buxbaum, 2017) or the laws that constrain object movement (Fischer et al., 2016). In either case, the specificity of decoding typical tool grasps in hand-, rather than tool- and hand-, selective cortex challenges the popular interpretation that brain activation for viewing tool images is a reflection of sensorimotor processing linked to tool manipulation (Martin et al., 1996; Mahon et al., 2007; Fang & He, 2005; Grafton et al., 1997; Martin & Chao, 2001; also see Mahon & Caramazza, 2009).

Several differences between our study and previous research are worth discussing. First our univariate analysis revealed no relationship between mean activity and typicality, even in the IPS and LOTC regions where we observed multivariate effects. Previous studies have found greater univariate activation in occipito-temporal and/or fronto-parietal cortex for typical relative to atypical actions when participants viewed pictures and movies or pantomimed (Johnson-Frey et al., 2003; Valyear et al., 2010; Yoon et al., 2012; Mizelle et al., 2013; Przybylski & Króliczak, 2017). Our results appear to fit the claim that MVPA can reveal more fine-grained effects (Kriegeskorte, et al., 2006), as recently argued by Buchwald et al. (2018) when showing that pantomimed typical tool vs non-tool grasps could be decoded from a range of regions including premotor and intraparietal areas. We suspect that task differences are also an important contributing factor to the general lack of univariate effects. For example, our experiment involved fewer, less varied, exemplars than those used in these previous picture studies. Likewise, our grasp-to-touch paradigm is simpler than studies showing greater univariate activations in the left SMG, premotor cortex, LOTC and IPS when performing real tool-use actions (Brandi et al., 2014; Valyear et al., 2012) or haptically-guided typical tool grasps (Styrkowiec et al., 2019) relative to tool/non-tool control actions. Finally, in our study, grasping always involved a precision grip whereas previous studies employed power grasps which are better suited for certain actions with some specific tools. This factor may have led to the lack of typicality decoding effects in tool-selective cortex as these areas could be sensitive to both the side of the object being grasped and the function of particular grips (e.g., clenching; Buxbaum et al., 2006). We designed our precision grasping task in order to investigate tool affordances while carefully equating biomechanics between actions, such that decoding typicality was unlikely to be attributed to motor-related differences. Future real action studies manipulating the type of grasp (e.g., grasp vs use) are needed to further identify the content of information coded by visual hand-/tool-selective areas.

It is worth noting that we were unable to match the visual symmetry between object categories (our tools were asymmetric while the non-tools were symmetric) because asymmetric non-tool bars were perceived as tools by participants (i.e. the wider side perceived as a functional-end). Nonetheless, tool-specific decoding in hand-selective cortex is unlikely to be explained by simple effects of symmetry: if effects were related to symmetry comparable decoding effects should have been observed in symmetry-responsive regions (e.g., LOTC-Object; EVC; Beck, et al., 2006), particularly since they are also known to code motor-related information (e.g. Gallivan & Culham, 2015; Monaco et al., 2020).

In conclusion, parietal and occipital visual regions specialised for representing hands were found to encode information about the functional relationship between the grasping hand and a tool, implicating hand-selective cortex in motor control. These findings raise novel questions about the possibility that overlapping hand- and tool-selective regions are functionally distinct and begin to uncover which brain regions evolved to support tool-use, a defining feature of our species.

## Acknowledgements

We thank Jenna Green, Richard Greenwood, Holly Weaver, Iwona Szymura and Emmeline Mottram for support in data collection, Derek Quinlan for building the real action set-up, Stefania Bracci for sharing the visual localizer stimuli and Annie Warman for comments on draft manuscript. This work was funded by grant (184/14) from the BIAL Foundation awarded to S. Rossit & F.W. Smith.

